# Engagement-Dependent Neural Entrainment Underlies Dissociable tACS Effects on Item and Sequence Working Memory

**DOI:** 10.64898/2026.05.27.728324

**Authors:** Liang Shi, Gui Xue

## Abstract

A central challenge in neuromodulation research is to elucidate how exogenous electrical stimulation interacts with endogenous neural dynamics to shape cognitive function. Drawing on empirical evidence for dissociable oscillatory mechanisms underpinning item versus sequence working memory (WM), we employed transcranial alternating current stimulation (tACS) and magnetoencephalography (MEG) to establish a causal separation and characterize the underlying neurophysiological basis. We observed a clear behavioral and neural dissociation: prefrontal 6-Hz theta stimulation selectively enhanced item WM, whereas frontoparietal in-phase theta stimulation specifically improved sequence WM. MEG further revealed task-specific modulation effects: during sequence WM, where frontoparietal theta synchrony predicted memory success, in-phase theta stimulation selectively amplified inter-regional synchronization. In contrast, during item WM, where local theta power was indicative of memory success, the identical stimulation overall exerted no significant modulation effect. Critically, across both tasks and oscillatory measures (inter-regional synchronization and theta power), the magnitude of stimulation-induced modulation scaled with each region’s or connection’s contribution to memory performance, indicating that tACS proportionately amplifies actively engaged oscillatory processes. This engagement-dependent principle explains task- and region-specific neurophysiological effects and the variability in behavioral outcomes, providing a framework for designing precise, functionally aligned stimulation protocols to enhance translational efficacy.

## Introduction

Noninvasive brain stimulation serves as a potent tool to modulate neural activity and probe the causal mechanisms of cognition. However, stimulation outcomes remain highly variable across studies, tasks, and individuals (Booth et al., 2022; Imburgio & Orr, 2018; López-Alonso et al., 2014; Wiethoff et al., 2014). In particular, the efficacy of transcranial alternating current stimulation (tACS)—a method designed to interact with and entrain endogenous neural rhythms (Antal & Paulus, 2013)—can vary significantly even when delivered at the same frequency or targeting identical cortical regions (Alekseichuk et al., 2017; Brauer et al., 2018; Kleinert et al., 2017; Rostami et al., 2021). Evidence from domains such as selective attention, language processing, and large-scale cortical physiology indicates that stimulation effects can be region- and task-specific (Castrillon et al., 2020; Heinen et al., 2014; Meinzer et al., 2012). This inconsistency highlights a fundamental theoretical challenge: how do externally applied oscillatory fields interface with the brain’s ongoing, task-dependent neural dynamics to influence behavior? While state-dependent accounts propose that stimulation interacts with the current functional state of the neural system (Alagapan et al., 2016; Bradley et al., 2022), the field still lacks a mechanistic framework that explains why identical stimulation parameters often produce variable effects across different tasks, individuals, and contexts. Specifically, it remains unknown which neural processes are preferentially modulated by tACS and how this modulation translates into task-specific behavioral outcomes. Addressing this gap is the central aim of the present study.

Theta-band activity serves as an important test case for such a framework. Theta oscillations are widely involved in working memory (WM), with frontal theta power associated with maintaining item-specific content (Otstavnov et al., 2024; Proskovec et al., 2018) and theta phase synchronization supporting the coordination of distributed WM networks (Payne & Kounios, 2009; Sarnthein et al., 1998; Sauseng et al., 2010; Wu et al., 2007). Recent evidence suggests that item and sequence WM may exert differential demands on these theta mechanisms, even when recruiting overlapping frontoparietal regions (Attout et al., 2019; Marshuetz et al., 2000; Rottschy et al., 2012). Specifically, item WM has been linked to increases in local theta power (Brookes et al., 2011; Fernández et al., 2021; Jensen & Tesche, 2002; Maurer et al., 2015), whereas sequence WM imposes additional requirements for maintaining temporal or ordinal structure and may rely more heavily on frontoparietal or hippocampal–prefrontal theta coupling (Chander et al., 2016; Johnson et al., 2018; Yang et al., 2021). Direct comparisons indicate that local theta and inter-regional theta phase synchronization contribute partially dissociable to item versus sequence WM maintenance (Su et al., 2024), though the causal roles of these mechanisms remain to be determined.

tACS presents an opportunity to test these causal contributions by selectively enhancing local oscillatory power (e.g., through unifocal theta stimulation) or promoting inter-regional synchrony (e.g., via bifocal in-phase stimulation). Indeed, unifocal theta tACS applied to the parietal or frontal cortex has been shown to enhance item WM performance and increase theta power (Jaušovec et al., 2014; Pahor & Jaušovec, 2018; Wolinski et al., 2018). In contrast, bifocal in-phase theta stimulation can strengthen frontoparietal coherence and facilitate WM retention (Hu et al., 2022; Polanía et al., 2012), whereas anti-phase stimulation often disrupts performance and reduces connectivity (Alekseichuk et al., 2017; Biel et al., 2022). Nevertheless, no existing studies have examined the dissociated behavioral effects and the underlying neurophysiological mechanisms of unifocal and bifocal stimulation for item and sequence WM, which may aid in developing a unified framework for accounting for these task- and region-specific effects.

The present study aims to establish such a framework in the context of WM. In addition to identifying the dissociation of item and sequence WM, we utilized their partially differentiable oscillatory signatures as a natural testbed for examining how tACS interacts with task-engaged neural processes. In two controlled tACS experiments, we applied unifocal prefrontal theta stimulation and bifocal frontoparietal in-phase or anti-phase theta stimulation while participants performed item and sequence WM tasks. In experiment 3, we employed magnetoencephalography (MEG) to quantify how in-phase theta stimulation modulates local theta power and frontoparietal theta phase synchronization during item and sequence WM maintenance. This design allowed us to test three hypotheses. First, if item WM relies primarily on local prefrontal theta power, unifocal theta tACS should selectively enhance item WM, whereas bifocal in phase theta tACS—targeting frontoparietal synchronization—should selectively enhance sequence WM. Second, if tACS interacts with task engaged processes, the same in phase stimulation should increase frontoparietal theta synchronization only during sequence WM, the task where this mechanism is behaviorally relevant, and not during item WM. Third, and most critically, we predicted an engagement dependent principle: the magnitude of tACS induced neural modulation should scale with each neural process’s contribution to memory success, as indexed by the subsequent memory effect. Our results provide a rigorous test of this principle, showing that externally applied theta fields preferentially and proportionally enhance the oscillatory processes actively supporting ongoing cognition. This framework moves beyond merely demonstrating causal effects to explaining their variability and context dependence, offering a mechanistic foundation for designing precise, functionally aligned stimulation protocols.

## Results

In this study, we conducted three tACS experiments to investigate how exogenous stimulation interacts with and modulates endogenous neural dynamics to influence cognition, particularly within the context of WM (Fig. 1A). Experiment 1 enrolled 39 participants, who were randomly assigned to either the unifocal stimulation group (n=20) or the bifocal stimulation group (n=19). Each participant completed three counterbalanced experimental conditions across separate test days: the unifocal group experienced 6 Hz stimulation, 35 Hz stimulation, and sham stimulation, while the bifocal group received 6 Hz in-phase stimulation, 6 Hz anti-phase stimulation, and sham stimulation (Fig. 1C). Stimulation frequency was set to 6 Hz, a central theta frequency (4–8 Hz) known to support WM maintenance (Jensen & Tesche, 2002; Sauseng et al., 2005) and previously shown to be effective in tACS studies (Jaušovec et al., 2014; Pahor & Jaušovec, 2018; Wolinski et al., 2018). During the 20-minute tACS sessions, followed by a 40-minute post-stimulation phase, all participants performed four WM tasks, including two item WM tasks and two sequence WM tasks (Fig. 1B). Both task types utilized the same two stimulus categories: fractal images and spatial locations, which allowed for clear and generalizable contrasts between item and sequence WM. To enhance the sensitivity of the stimulation effects, task difficulty was individually calibrated prior to the formal experiment using an adaptive procedure (i.e., one-up-two-down method). For both item and sequence WM tasks, participants initially viewed a series of sequentially presented stimuli (either fractal images or spatial locations). Following a 2-second delay, they were required either to select the presented stimuli from a set of distractors (item WM) or accurately reconstruct the order of the presented stimuli (sequence WM).

**Fig. 1.**
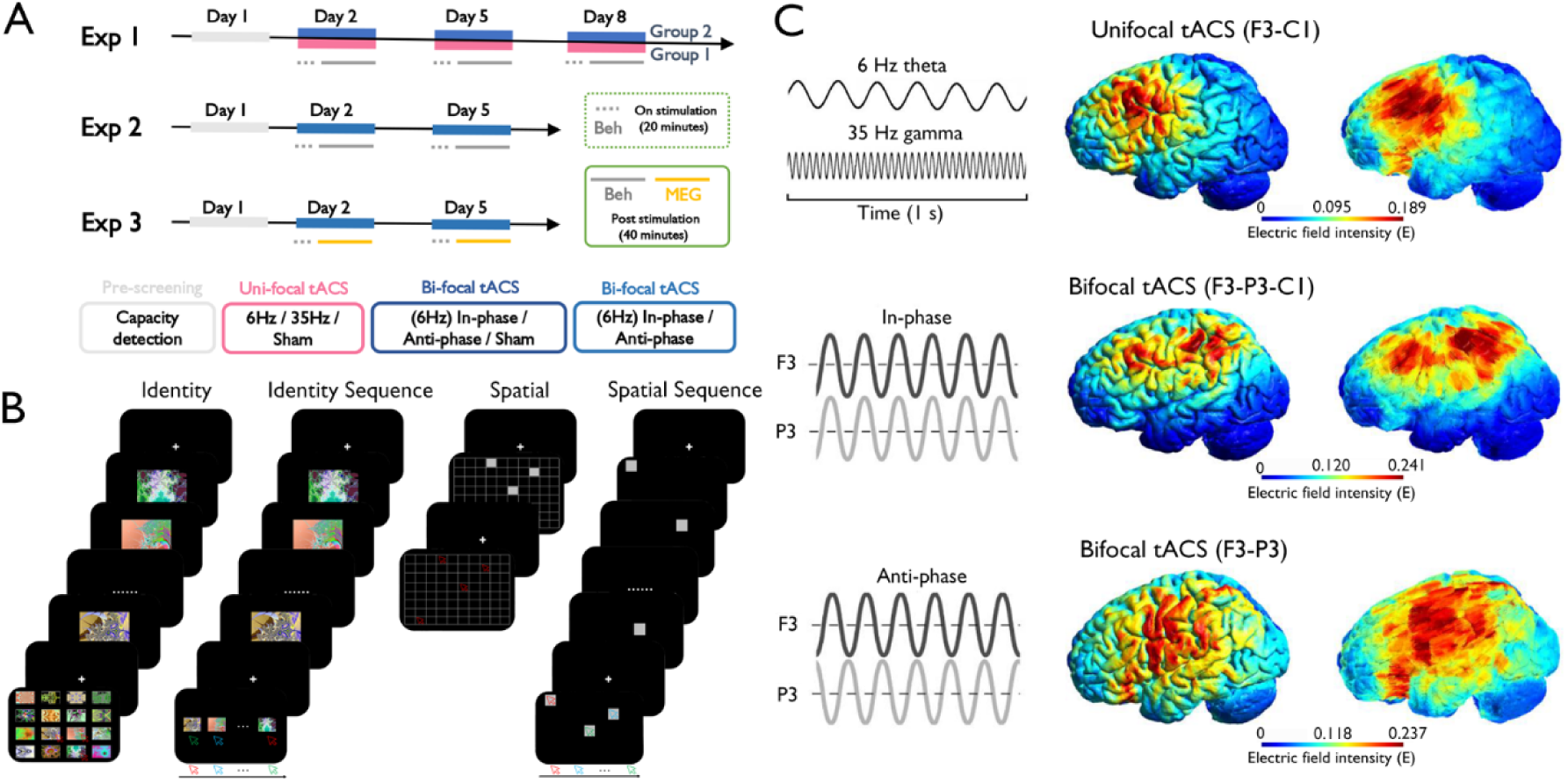
Experimental Design and tACS Protocols. **(A) Overview of Experimental Design.** This figure illustrates the structure of three randomized, single-blind experiments. Experiment 1 encompassed four days, including one pre-screening session and three transcranial alternating current stimulation (tACS) sessions. A between-group design was employed: Group 1 received unifocal tACS (6 Hz, 35 Hz, or sham) over the left prefrontal cortex (F3) on different days, while Group 2 underwent bifocal tACS (6 Hz in-phase, 6 Hz anti-phase, or sham) targeting the frontoparietal network (F3–P3). Experiments 2 and 3, lasting three days each, incorporated one pre-screening session and two tACS sessions, focusing solely on bifocal 6 Hz tACS (both in-phase and anti-phase). Additionally, Experiment 3 included a post-stimulation magnetoencephalography (MEG) session. (B) **Memory Tasks.** Four tasks were administered across all experiments to assess item working memory (WM) (recognizing previously encoded fractal images or spatial locations) and sequence WM (reconstructing the order of encoded fractal images or spatial locations). (C) **tACS Protocols.** The top panel displays unifocal tACS (6 Hz or 35 Hz) targeted at F3, with a return electrode placed at C1 (stimulation intensity: 2 mA). The middle panel illustrates bifocal in-phase 6 Hz tACS (stimulation intensity: 1 mA at F3 and P3, and 2 mA at C1). The bottom panel depicts bifocal anti-phase 6 Hz tACS (180° phase difference between F3 and P3), with stimulation intensity set at 1 mA for both F3 and P3. The corresponding cortical E-field magnitude distributions for each protocol are also visualized

### Dissociable behavioral effects on item and sequence WM

Experiment 1 investigated the differential effects of unifocal and bifocal tACS on item and sequence WM, building on the partial dissociation of local versus inter-regional theta mechanisms (Su et al., 2024). In the unifocal tACS group (F3-targeted), a 2 (WM task: item vs. sequence) × 2 (stimulus: fractal vs. spatial) × 3 (stimulation condition: 6 Hz vs. 35 Hz vs. sham) ANOVA revealed a significant interaction between WM task and stimulation condition (*F_1.945, 35.021_ = 3.793, p = 0.033, partial eta-squared* (η*_p_^2^) = 0.174*). No other significant main effects or interactions were identified (all *Ps* > 0.118). Subsequent analyses indicated a significant main effect of stimulation condition for the item WM task (*F_1.698, 30.578_ = 4.247, p = 0.029*, *η_p_^2^ = 0.104*), indicating superior WM performance under 6 Hz stimulation compared to both 35 Hz (*t_18_ = 2.823, p = 0.011, Cohen’s d = 0.541*) and sham (*t_18_ = 2.577, p = 0.019, Cohen’s d = 0.437*) conditions (Fig. 2A). No significant effects were observed for the sequence WM task (all *Ps* > 0.106).

For the bifocal tACS group (F3-P3-targeted), a 2 (WM task: item vs. sequence) × 2 (stimulus: fractal vs. spatial) × 3 (stimulation condition: in-phase vs. anti-phase vs. sham) ANOVA also yielded a significant interaction between WM task and stimulation condition (*F_1.952, 30.086_ = 5.049, p = 0.021*, η*_p_^2^ = 0.210*). Subsequent analyses indicated a significant main effect for the sequence WM task (*F_1.582, 30.050_ = 5.333, p = 0.015*, η*_p_^2^ = 0.219*), with superior WM performance observed under in-phase stimulation compared to anti-phase (*t_19_ = 2.794, p = 0.012, Cohen’s d = 0.492*) and sham (*t_19_ = 3.391, p = 0.003, Cohen’s d = 0.354*) conditions (Fig. 2B). No significant effects were found for the item WM task (all *Ps* > 0.192).

The double dissociation between item and sequence WM was further confirmed when examining the stimulation-induced performance changes (i.e., 6Hz – 35 Hz for unifocal stimulation and in-phase – anti-phase for bifocal stimulation as capacity gain). A 2 (WM task: item vs. sequence) × 2 (stimulus: fractal vs. spatial) × 2 (stimulation type: unifocal vs. bifocal) mixed-design ANOVA revealed a significant interaction between WM task and stimulation type (*F_1, 37_ = 9.728, p = 0.004,* η*_p_^2^ = 0.208*). Post-hoc tests confirmed that unifocal stimulation induced greater improvement in item as compared to sequence WM (*F_1, 18_ = 8.655, p = 0.009,* η_*p*_^2^ = 0.325); *t_18_ = 2.942, p = 0.009, Cohen’s d = 0.668*), whereas a reversed pattern was evident for bi-focal stimulation (*F_1, 19_ = 4.834, p = 0.040,* η*_p_^2^ = 0.115; t_19_ = 2.194, p = 0.040, Cohen’s d = 0.366)*.

In short, Experiment 1 demonstrated that prefrontal theta stimulation selectively enhances item WM performance, while frontoparietal in-phase theta stimulation preferentially improves sequence WM performance.

**Fig. 2.**
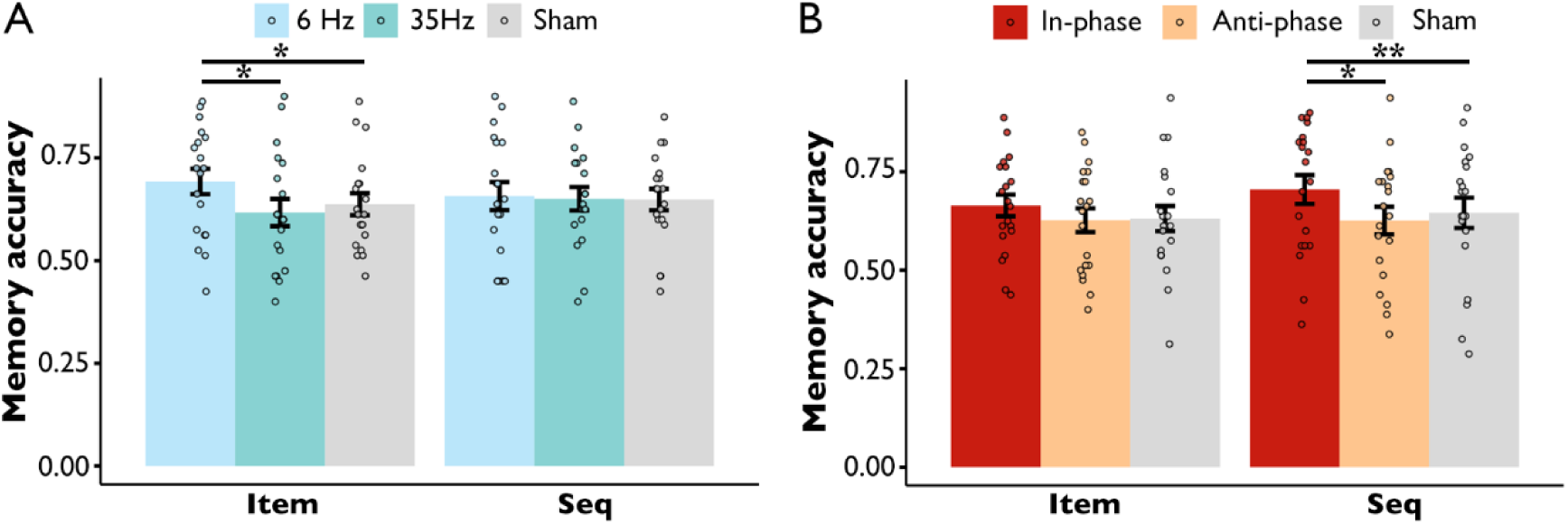
Dissociable Effects of tACS on Item and Sequence Working Memory (WM) in Experiment 1. (A) **Unifocal Prefrontal tACS.** Theta (6 Hz) stimulation over the left prefrontal cortex (F3) significantly enhanced item WM accuracy compared to both 35 Hz stimulation and sham conditions, while no significant effect was noted on sequence WM. (B) **Bifocal Frontoparietal tACS.** In-phase theta (6 Hz) stimulation targeting the F3–P3 network specifically improved sequence WM performance relative to anti-phase and sham conditions. Bifocal tACS did not demonstrate significant effects on item WM. \**p* < 0.05, \*\**p* < 0.01.

### Replication of sequence WM enhancement by frontoparietal in-phase theta tACS using MEG

Having identifying the specific role of frontoparietal in-phase theta stimulation in enhancing sequence WM, we utilized MEG to further explore the underlying neural mechanisms. Given that only magnet-compatible keyboards could be utilized for behavioral responses in the MEG scanner, the behavioral paradigm was slightly modified (Fig. S1). Prior to the MEG study (Experiment 3, N = 23), a behavioral experiment (Experiment 2, N = 26) was conducted to replicate the stimulation effect. Only 6 Hz in-phase and anti-phase stimulations were included, as Experiment 1 revealed no significant differences between anti-phase and sham conditions.

An experiment (Experiment 2 vs. Experiment 3) × WM task (item vs. sequence) × stimulus (fractal vs. spatial) three-way ANOVA on memory gain (calculated as in-phase – anti-phase) showed no main effect of experiment (*F_1, 47_ = 0.061, p = 0.805,* η*_p_^2^* = 0001) or any interactions involving experiment (all *Ps* > 0.660). We then pooled data from both experiments and subjected them to a 3-way ANOVA (WM task × stimulus × stimulation condition: in-phase vs. anti-phase), which revealed a significant interaction between WM task and stimulation condition (*F_1, 48_ = 7.308, p = 0.010,* η*_p_^2^ = 0.132*) and a significant main effect of stimulation condition (*F_1, 48_ = 9.931, p = 0.003,* η*_p_^2^ = 0.171*). No other main effects or interactions were significant (all *Ps* > 0.162).

For the sequence WM task, a 2-way ANOVA (stimulus × stimulation condition) showed a significant main effect of stimulation condition (*F_1, 48_ = 24.212, p < 0.001,* η*p^2^ = 0.335*), indicating superior WM performance under in-phase compared to anti-phase stimulation (Fig. 3); no main effects of stimulus or interactions were observed (all *Ps* > 0.839). In contrast, for the item WM task, no significant main effects or interactions were found (all *Ps* > 0.135). These patterns were replicated separately for Experiment 2 (sequence WM: *F_1, 25_ = 13.124, p = 0.001,* η*_p_^2^ = 0.344*; item WM: *F_1, 25_ = 0.419, p = 0.523,* η*_p_^2^ = 0.017*) and Experiment 3 (sequence WM: *F_1, 22_ = 11.125, p = 0.003,* η*_p_^2^ = 0.336*; item WM: *F_1,22_ = 0.956, p = 0.338,* η*_p_^2^ = 0.045*).

**Fig. 3.**
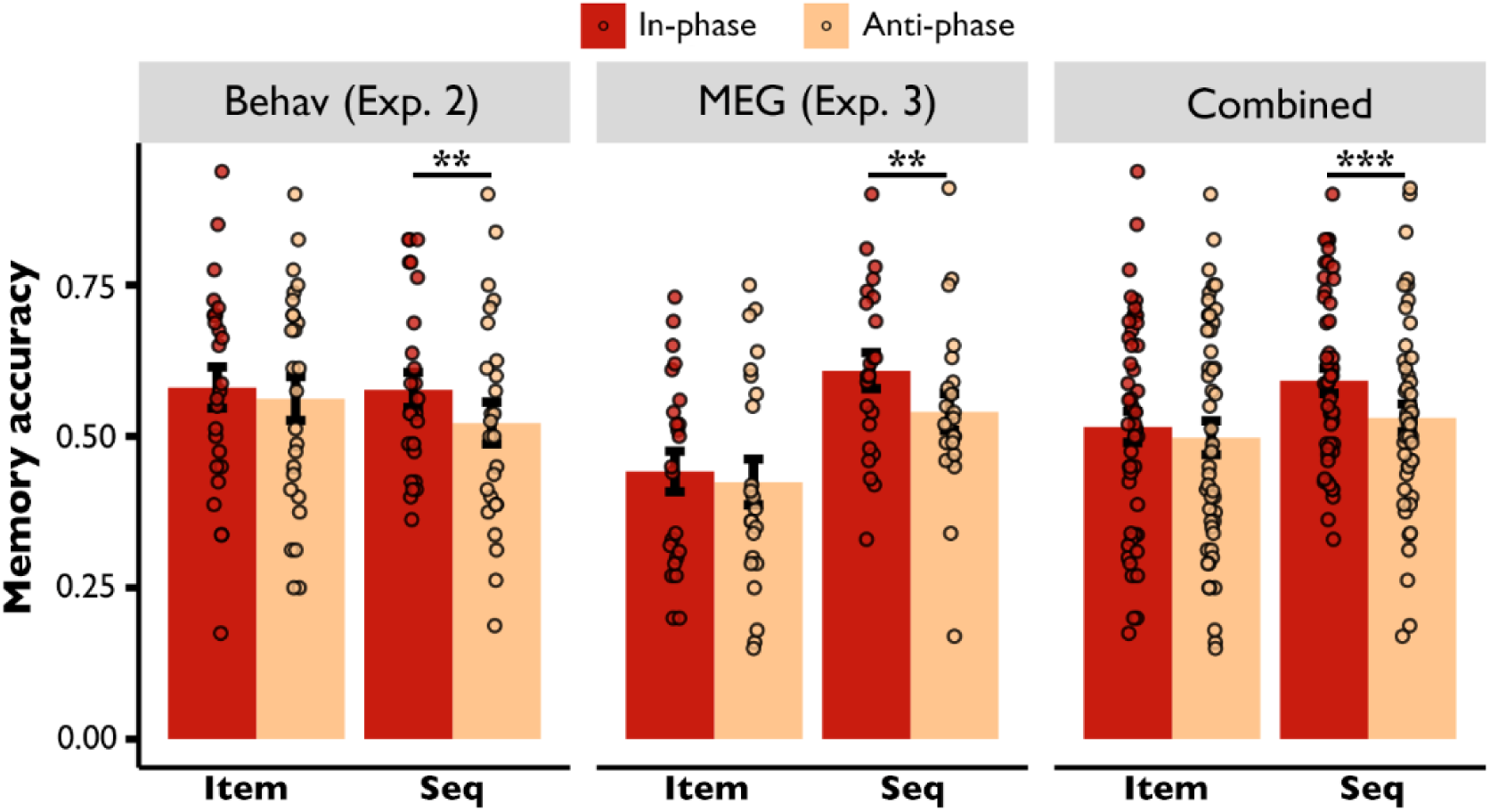
Replicable Enhancement of Sequence Working Memory (WM) via Frontoparietal In-Phase Theta tACS in Experiments 2 and 3. In both experiments, in-phase theta (6 Hz) stimulation over the F3–P3 network significantly enhanced accuracy for sequence WM compared to anti-phase stimulation. This effect was replicated despite modifications in response methods (numeric keypad) adapted for different testing environments: a behavioral room (Experiment 2) and an MEG scanner (Experiment 3). No significant differences were observed between stimulation conditions for the item WM task. \*\**p* < 0.01, \*\*\**p* < 0.001.

### Frontoparietal theta phase synchronization contributed to sequence WM maintenance

To understand the neural mechanisms underlying the dissociated stimulation effect, we first examined the neural substrates of successful sequence WM maintenance. To quantify phase-based functional connectivity, the debiased weighted phase-lag index (dwPLI) (Vinck et al., 2011) was calculated for each channel pair. We analyzed the subsequent memory effect (SME)—defined as significantly higher dwPLI values for remembered versus forgotten trials in the sequence WM task—using only data from the anti-phase stimulation condition to exclude tACS-induced interference.

To identify which edges showed significant SME, we examined all 36 channel pairs by conducting WM task (item vs. sequence) × stimulus (fractal vs. spatial) × SME (remembered vs. forgotten) ANOVAs. Two edges survived FDR correction, which showed significant WM task × SME interactions and no significant three-way interactions (*p > 0.468*; Fig. 4A). These edges included: left frontal-left parietal (LF-LP, *F_1, 22_ = 13.032, p = 0.002, p_fdr_ = 0.028,* η*p^2^ = 0.268*) and left fontal-left occipital (LF-LO, *F_1, 22_ = 14.969, p < 0.001, p_fdr_ = 0.028,* η*p^2^ = 0.405*; Fig. 4B). Subsequent 2-way ANOVAs (stimulus × SME) confirmed that SMEs were exclusive to the sequence WM task (*p < 0.005,* η*p^2^ > 0.306* for SME, *p > 0.120* for interaction) and absent in the item WM task (*p > 0.218* for SME, *p > 0.065* for interaction), indicating these edges specifically support sequence WM maintenance.

### In-phase theta stimulation selectively increased frontoparietal theta phase synchronization during sequence WM maintenance

Next, to examine the effect of frontoparietal in-phase theta stimulation on phase synchronization within sequence WM-related edges, we conducted three-way ANOVAs with factors of WM task (item vs. sequence) * stimulus (fractal vs. spatial) * stimulation condition (in-phase vs. anti-phase) on the above FDR-corrected edges. Only the LF-LP edge exhibited a significant interaction between WM task and stimulation condition (*F_1, 22_ = 8.035, p = 0.010, p_fdr_ = 0.019,* η*p^2^ = 0.268*), with no significant three-way interaction (*F_1, 22_ = 0. 112, p = 0.741,* η*p^2^ = 0.005*). Subsequent 2-way ANOVAs (stimulus × stimulation condition) conducted separately for each WM task demonstrated a significant tACS effect specific to the sequence WM task at the LF-LP edge (main effect of stimulation condition: *F_1, 22_ = 4.670, p = 0.041,* η*p^2^ = 0.176*; interaction: *F_1, 22_ = 0.239, p = 0.630,* η*p^2^ = 0.011*). No significant modulation was detected for the item WM task (main effect of stimulation condition: *F_1, 22_ = 2.544, p = 0.125,* η*_p_^2^ = 0.104*; interaction: *F_1, 22_ = 0.013, p = 0.909,* η*_p_^2^ = 0.001*). Post-hoc comparisons confirmed that in-phase theta stimulation significantly enhanced LF-LP synchronization during sequence WM task compared to anti-phase stimulation (*t_22_ = 2.168, p = 0.041, Cohen’s d = 0.344*; Fig. 4C). No other edges exhibited significant tACS modulation in either sequence WM (all *Ps* > 0.063) or item WM tasks (all *Ps* > 0.107).

If LF-LP synchronization contributes to the behavioral effects of tACS, individuals demonstrating greater enhancement of LF-LP synchronization should exhibit larger behavioral improvements. A significant positive correlation was observed between LF-LP dwPLI gain (in-phase minus anti-phase) and sequence WM behavioral gain using robust correlation (r = 0.454, p = 0.029, 95% CI = [0.052 0.730]). No such correlation was found for the item WM task (r = -0.078, p = 0.723, 95% CI = [-0.038 0.684]). A Fisher z-test comparing the two correlation coefficients revealed a significant difference (z = 1.750, p = 0.040).

Collectively, these findings provide converging evidence that LF-LP synchronization is specifically involved in sequence WM, and its enhancement mediates the behavioral improvements induced by frontoparietal in-phase theta stimulation. The results demonstrate selective modulation of frontoparietal connectivity by theta tACS, with effects that are task-and edge-specific.

**Fig. 4.**
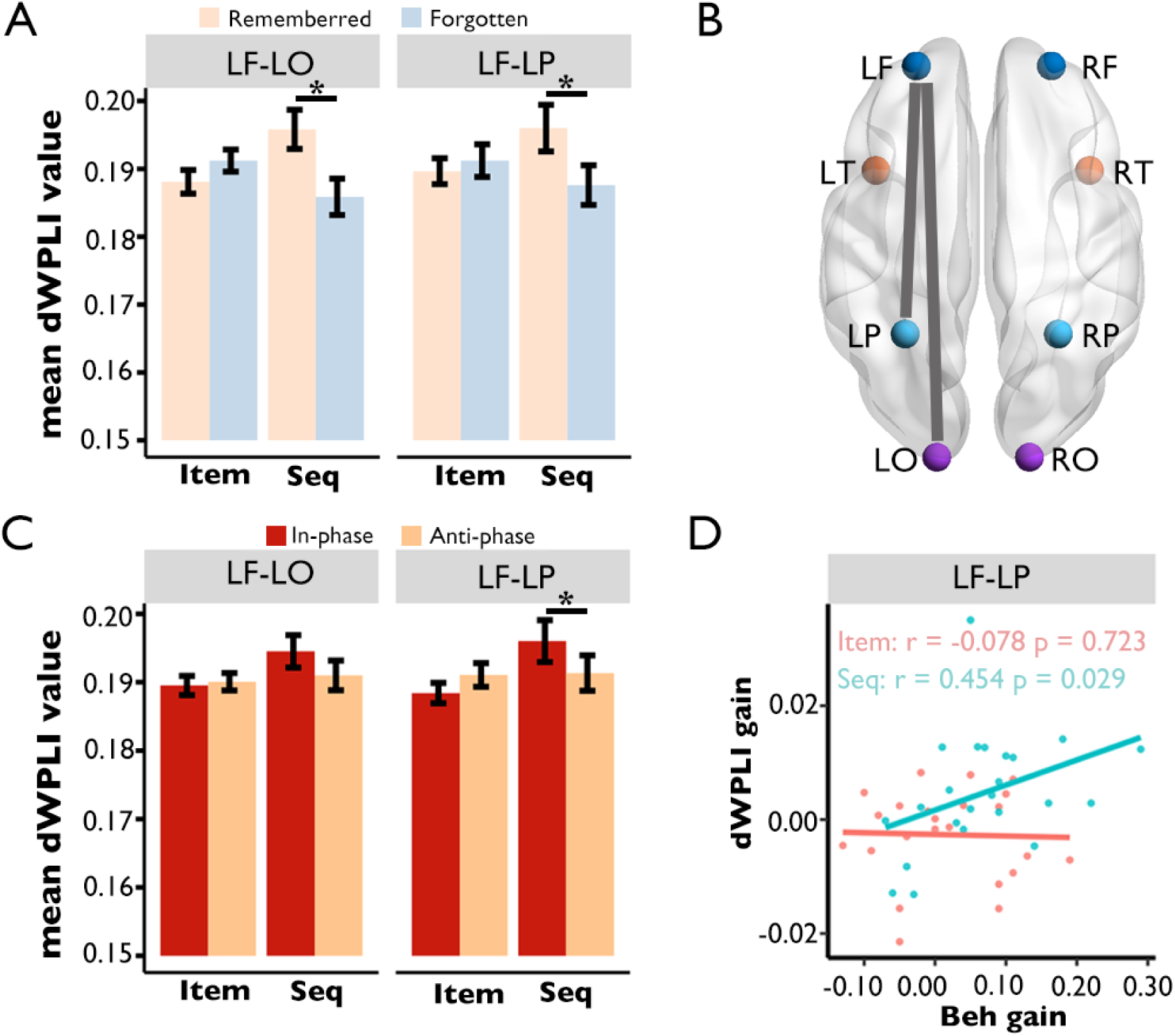
Subsequent Memory Effects (SMEs) and Task-Specific Connectivity Modulations Quantified via Debiased Weighted Phase-Lag Index (dwPLI). (A) Mean dwPLI values for two brain edges were significantly higher for remembered than forgotten trials in the sequence WM task, reflecting a robust SME. (B) Schematic representation of the two edges exhibiting significant SME in sequence WM: left frontal-left parietal (LF-LP) and left frontal-left occipital (LF-LO). (C) Modulatory effects of tACS on SME-related edges: only the LF-LP edge demonstrated significantly enhanced synchronization under in-phase compared to anti-phase stimulation during sequence WM, with no tACS effects observed on the LF-LO edge. (D) robust correlation analysis: a significant positive correlation was identified between LF-LP dwPLI gains (in-phase – anti-phase) and sequence WM capacity gains, whereas no such correlation was present for the item WM task. \**p* < 0.05.

### The tACS effect on phase synchronization was modulated by task involvement

The above results demonstrate task-specific tACS effects: frontoparietal 6 Hz in-phase tACS selectively enhanced theta phase synchronization during sequence WM task but not during item WM task. A plausible explanation is that this modulation is contingent upon task-specific neural engagement. To test this hypothesis, we computed the dwPLI SME for each edge and task type.

On average, the effect size of SME on dwPLI was significantly weaker for item WM compared to sequence WM tasks (*t_35_ = -4.975, p < 0.001, Cohen’s d = 1.258*, aligning with the specialized role of theta phase synchronization in sequence WM. Additionally, the mean effect size of tACS (in-phase vs. anti-phase) on dwPLI was more pronounced for sequence WM than for item WM (*t_35_ = 5.389, p < 0.001, Cohen’s d = 1.344*), further corroborating the specificity of in-phase theta stimulation in enhancing theta phase synchronization for sequence WM.

Intriguingly, a significant positive correlation was observed between the effect size of SME and tACS across edges for the sequence WM task (*r = 0.520, p = 0.001, 95% CI = [0.231 0.725]*), but not for the item WM task (*r = 0.182, p = 0.288, 95% CI = [-0.156 0.482]*). The latter might be due to overall weak stimulation effect and SME. To examine whether this engagement-dependent principle could account for the results in both tasks, we did the same correlation analysis by pooling together from both tasks. Indeed, we found a significant correlation between the effect size of SME and tACS (*r = 0.505, p < 0.001, 95% CI = [0.310 0.660]*; Fig. 5).

Collectively, these findings provide strong evidence that in-phase theta tACS enhances sequence WM by selectively modulating phase synchronization within the task-relevant neural circuitry, and this engagement-dependent effect provide a unified account for the task- and edge-specific modulation effect.

**Fig. 5.**
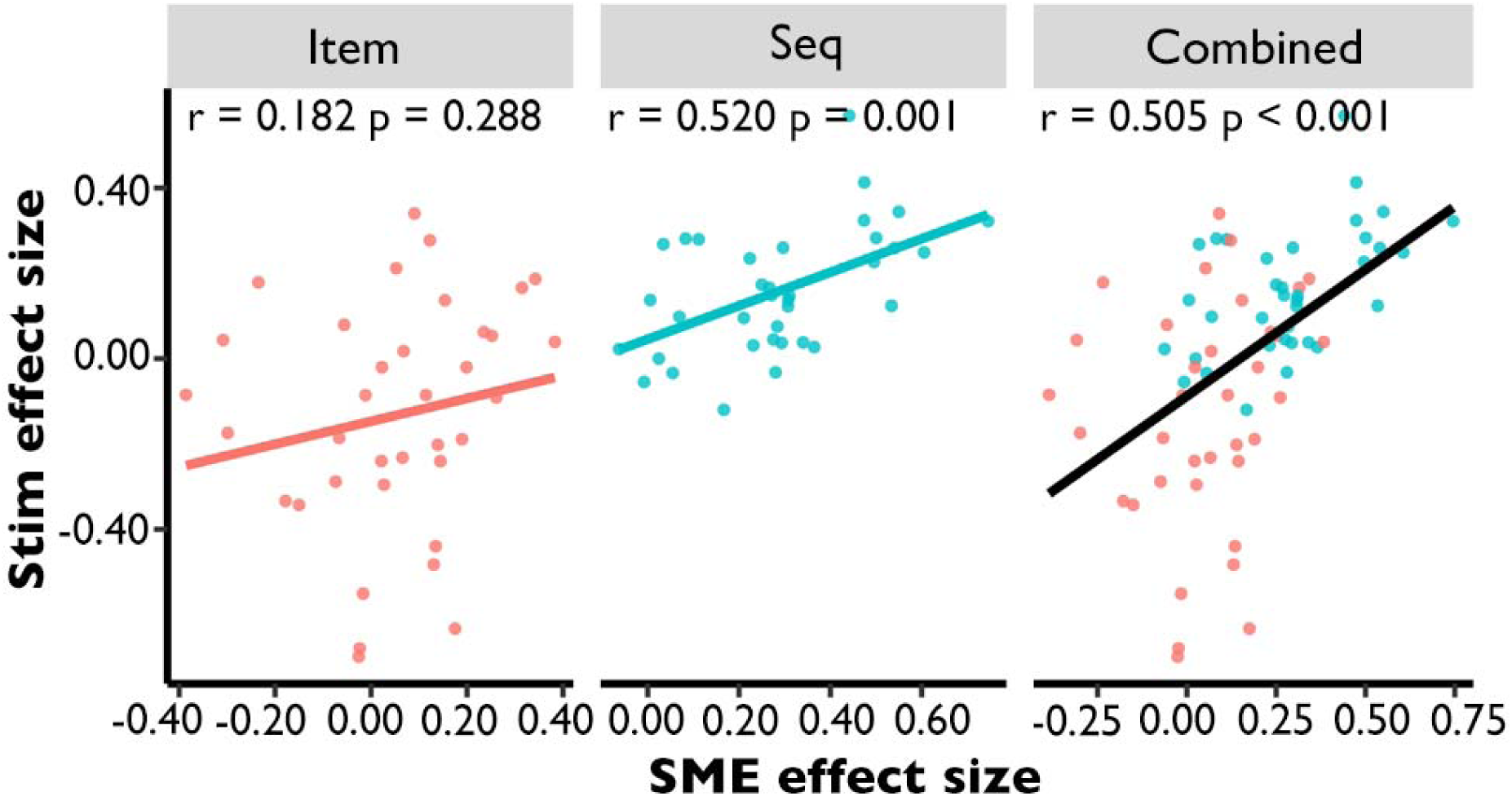
Correlation Between Subsequent Memory Effect (SME) and tACS Effect Sizes on Inter-Regional Theta Phase Synchronization. This scatterplot depicts the relationship between SME effect size (quantifying task-specific neural engagement) and tACS effect size (in-phase vs. anti-phase) for inter-regional theta phase synchronization. A significant positive correlation was found for the sequence WM task and when data from sequence and item WM tasks were pooled. In contrast, no significant correlation was detected for the item WM task.

### Frequency match between stimulation and endogenous theta frequency predicts the effect of neural entrainment

Our engagement dependent principle demonstrates that tACS effects scale with task relevance across different neural processes within each participant. Meanwhile, a separate, but not competing prediction from the resonance/frequency-match hypothesis is that the efficacy of tACS should also depend on the match between the stimulation frequency and each individual’s endogenous oscillatory frequency (Krause et al., 2022). To test this hypothesis, we estimated each participant’s task engaged theta peak frequency during the anti phase stimulation stage and examined whether the mismatch with the fixed 6 Hz stimulation predicts the magnitude of tACS induced neural and behavioral effects.

We found no significant correlation between the frequency mismatch and behavioral gains (item WM: r = − 0.327, p = 0.128, 95% CI = [-0.667 0.010]; sequence WM: r = −0.125, p= 0.569, 95% CI = [-0.497 0.286]; Fig. 6A). However, for the sequence WM task, there was a significant negative correlation between mismatch and dwPLI gain (r = −0.421, p = 0.046, 95% CI = [-0.719 -0.004]; Fig. 6B), indicating that participants whose endogenous theta frequency was closer to 6 Hz exhibited larger tACS-induced increases in frontoparietal theta synchronization. Importantly, this relationship was not observed for the item WM task (r = 0.021, p = 0.925, 95% CI = [-0.413 0.452]). Fisher z-test comparing the two correlation coefficients revealed approached significance (z = 1.448, p = 0.074). Together, this interaction supports an engagement-dependent resonance model: tACS benefits from individual frequency tuning only when the targeted neural process is actively recruited by the task.

**Fig. 6.**
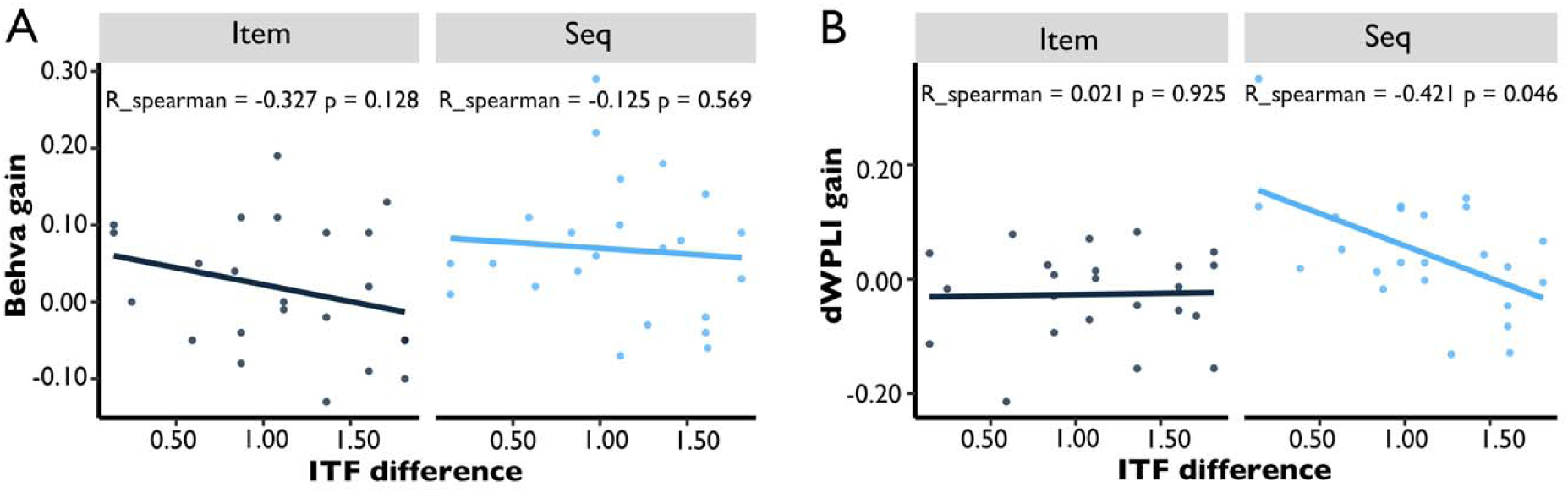
Correlation between Frequency Mismatch and the Magnitude of tACS Effect. (A) The scatter plot showing no significant correlations between mismatch (individual time-frequency difference, ITF difference) and behavioral gains (accuracy difference: in-phase minus anti-phase) for both working memory (WM) tasks. (B) The scatter plot showing a negative correlation between mismatch and neural gains (dwPLI difference for the LF-LP edge, in-phase minus anti-phase) only for sequence WM task.

### Theta power contributed to item WM maintenance

Beyond examining inter-regional theta phase synchronization, we also investigated task-dependent differences in time-frequency power during the WM maintenance stage. To identify time frequency clusters with significant SME, we performed a whole brain cluster based permutation test. This exploratory analysis revealed an early alpha cluster and a late theta cluster in the item WM task (Fig. 7A): an early alpha cluster (peak frequency = 13 Hz; time window = 0.26–0.59 s; *p < 0.038*, cluster-corrected) and a late theta cluster (peak frequency = 5 Hz; time window = 1.64–2.00 s; *p < 0.043*, cluster-corrected). By contrast, no significant clusters were detected in the sequence WM task (*cluster-level p > 0.080*).

To demonstrate the functional specificity of these two clusters, we conducted two separate three-way ANOVAs, including WM task (item vs. sequence), stimulus (fractal vs. spatial), and SME (remembered vs. forgotten) as independent variables. For the early alpha cluster, we observed a significant main effect of SME (*F_1, 22_ = 5.091, p = 0.034,* η*p^2^ = 0.192*), whereas neither the WM task × SME interaction nor the three-way interaction reached statistical significance (all *Ps* > 0.109). In contrast, analysis of the late theta cluster yielded a significant WM task × SME interaction (*F_1, 22_ = 11.624, p = 0.003,* η*p^2^ = 0.346*), with no other significant two-way or three-way interactions (all *Ps* > 0.258; Fig. 7B). Post-hoc comparisons confirmed that in the item WM task, remembered trials were associated with lower theta power than forgotten trials (t_22_ = −3.419, p = 0.002, *Cohen’s d = 0.703*). A reversed pattern, although statistically nonsignificant (t_22_ = 1.392, p = 0.178, *Cohen’s d = 0.227*), was found for the sequence WM task.

To elucidate how frontoparietal in-phase theta stimulation modulate theta power within the late theta cluster, we performed an additional three-way repeated-measures ANOVA including WM task (item vs. sequence), stimulus (fractal vs. spatial), and stimulation condition (in-phase vs. anti-phase). This analysis revealed a significant WM task × stimulation condition interaction (*F_1, 22_ = 9.208, p = 0.007,* η*p^2^ = 0.295*), and the three-way interaction was nonsignificant (*F_1, 22_ = 0.443, p = 0.512,* η*p^2^ = 0.020*). Follow-up two-way ANOVAs (stimulus × stimulation condition), stratified by WM task, found overall nonsignificant main effects of stimulation condition or interactions in either the item WM (all *Ps* > 0.130) or sequence WM (all *Ps* > 0.109; Fig. 7C), a finding consistent with the absence of significant behavioral improvement in item WM performance. Notably, there was a trend toward decreased theta power for the item WM after in-phase stimulation (t_22_ = -0.367), whereas a trend toward increased theta power for the sequence WM (t_22_ = 1.670). These directional trends align with the negative theta SME observed in item WM and the trending positive SME observed in sequence WM.

In short, these results suggest that later theta power was specifically involved in item WM, and frontoparietal in-phase theta stimulation modulated task-relevant theta power in a manner mirroring the distinct functional contributions of theta oscillations to item versus sequence WM.

**Fig. 7.**
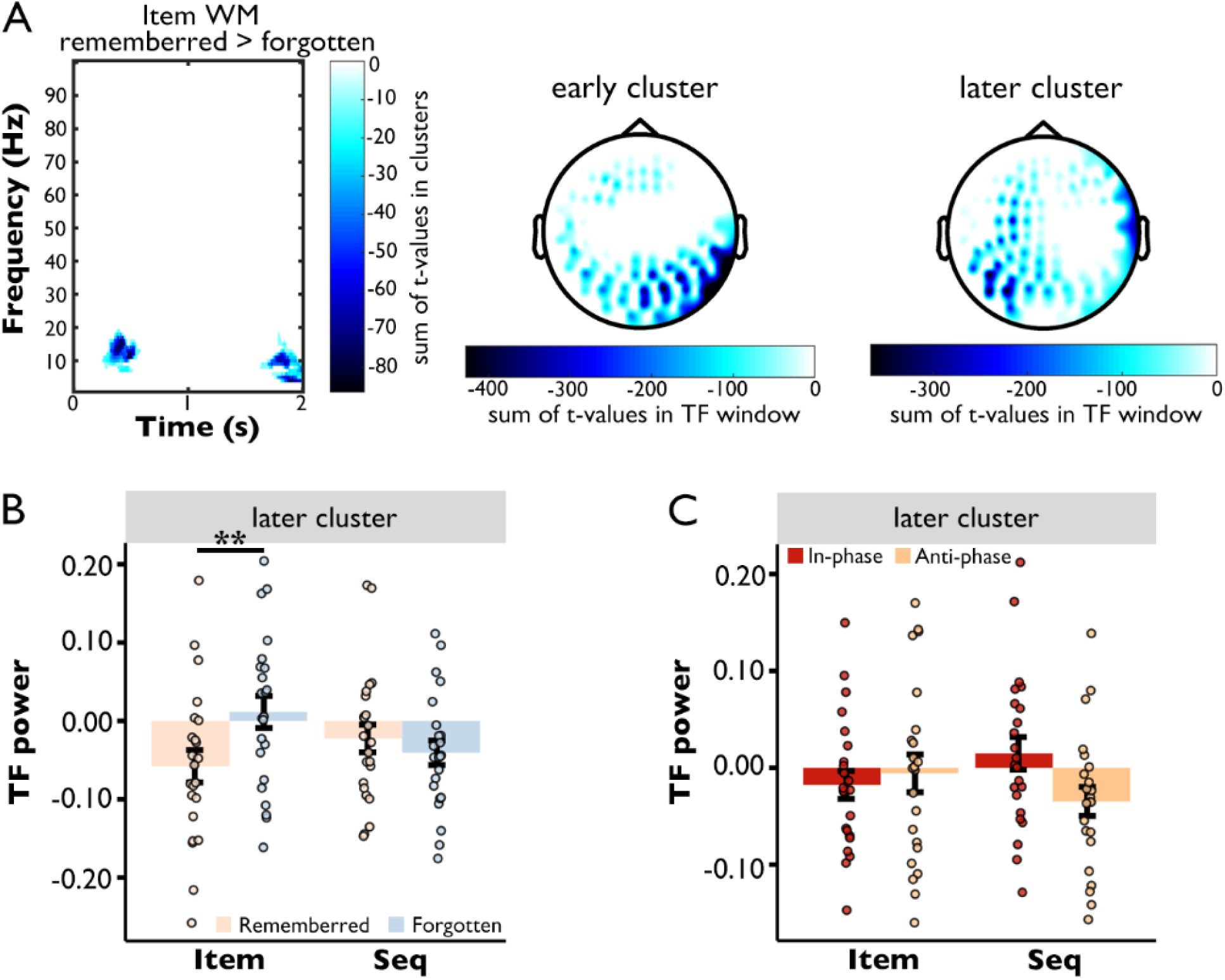
Theta Power Contributed to Item Working Memory (WM) Maintenance. (A) Left panel: Cluster-based permutation testing of the subsequent memory effect (SME) contrast (remembered vs. forgotten trials) identified two distinct clusters exhibiting significant SMEs in the item WM task. Middle and right panels: Grand-average topographic maps corresponding to these two significant clusters. (B) SME for late theta cluster power: In the item WM task, late theta cluster power was significantly lower for remembered than forgotten trials, with no such differences observed in sequence WM. (C) tACS modulation of late theta cluster power: No significant differences were detected between in-phase and anti-phase stimulation conditions for either item or sequence WM tasks. \*\**p* < 0.01.

### The tACS effect on theta power was modulated by task involvement

To further examine whether there was a region-specific stimulation effect for theta power that was determined by neural engagement, we also correlated the theta SME effect size with the tACS effect size (in-phase vs. anti-phase) across all MEG channels for each task. Results revealed a significant positive correlation for the item WM task (*r = 0.290, p < 0.001, 95% CI = [0.184 0.390]*), but not for the sequence WM task (*r = 0.093, p = 0.104, 95% CI = [-0.019 0.203]*). When data from both tasks were pooled, a significant correlation was also found (*r = 0.306, p < 0.001, 95% CI = [0.232 0.376]*; Fig. 8). Together, these results suggest that even when the overall modulation effect was nonsignificant, the variable effects across tasks and regions could still be accounted for by the engagement-dependent principle.

**Fig. 8.**
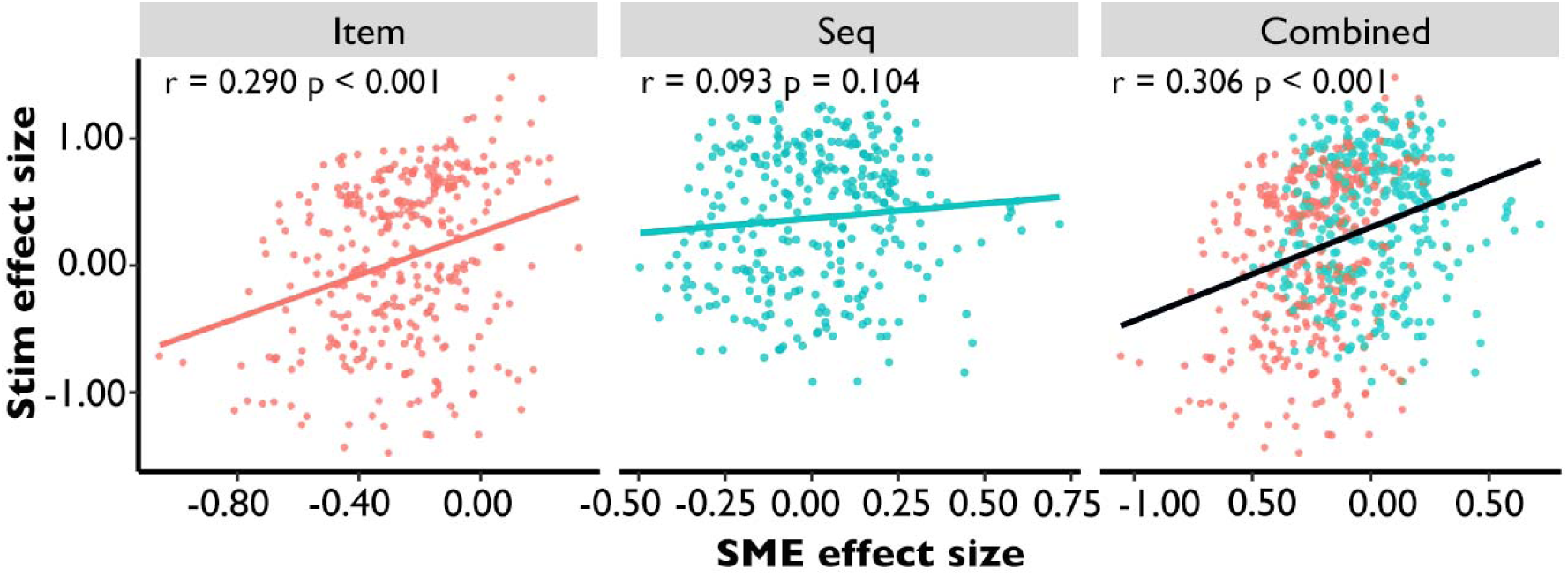
Correlation of Subsequent Memory Effect (SME) and tACS Effect Sizes on Local Theta Power. This scatterplot illustrates a significant positive correlation between SME and tACS (in-phase vs. anti-phase) effect sizes for the item WM task and pooled task data, with no correlation observed for the sequence WM task.

## Discussion

The present study tackles a central challenge in neuromodulation: understanding how externally applied rhythmic stimulation interacts with the brain’s ongoing, task-engaged activity to influence cognition. To explore this question, we combined tACS with behavioral assays and MEG to investigate how identical stimulation interacts with two WM tasks. These tasks activate overlapping frontoparietal regions but differ in their reliance on local versus network-level theta mechanisms. Across both tasks, we found convergent evidence for task-specific and engagement-dependent modulation of neural activity and behavior. These results provide an integrative framework that explains variability in stimulation outcomes and offers mechanistic insights into how external oscillatory fields interface with endogenous brain dynamics.

A substantial body of research has demonstrated that the effects of noninvasive stimulation depend on the brain’s current functional state (Alagapan et al., 2016; Bradley et al., 2022). Stimulation does not uniformly affect cognitive contexts but interacts differently with neural dynamics across various states, such as sleep phases (Li et al., 2017; Massimini et al., 2005), awareness (Gersner et al., 2011), or attentional engagement (Kamke et al., 2012). For instance, transcranial magnetic stimulation (TMS) applied to the frontal eye field preferentially modulates attended networks during selective attention (Heinen et al., 2014; Morishima et al., 2009) and high-frequency (20 Hz) repetitive TMS enhances memory-relevant networks during autobiographical retrieval (Warren et al., 2019). Our study extends these observations, demonstrating that stimulation outcomes can be significantly shaped by task requirements alone, even when stimuli and timing are held constant. Specifically, we found that identical frontoparietal in-phase tACS produced task-specific effects: stimulation selectively enhanced sequence WM performance and increased frontoparietal theta synchronization when such synchrony was behaviorally relevant, yet failed to exert a significant effect when local oscillatory activity supported item WM maintenance.

Critically, we also found the in-phase theta stimulation differently modulated theta power in the two WM tasks, in a manner consistent with theta contribution to item and sequence WM. These findings provide compelling evidence to suggest that tACS selectively modulates neural processes that are actively engaged by the current task, even when stimulation parameters are held constant.

Importantly, the present study also refines our understanding of regional specificity in neuromodulation. Previous research has shown that stimulation effects vary across cortical regions (Castrillon et al., 2020) and depend on functional connectivity strength within the targeted areas (Ezzyat et al., 2024; Kim et al., 2018; Solomon et al., 2018). Our results further extend these findings by showing that regional susceptibility is constrained by each region’s functional engagement during the task, quantified through the SME. Specifically, stimulation-induced changes in frontoparietal synchronization correlated with the regions’ synchronization SME, while changes in theta power correlated with the regions’ theta SME. This suggests that stimulation does not merely modulate the anatomical targets of the electric field but instead affects brain regions actively contributing to task performance.

By pooling data from two tasks, we reveal a critical, engagement-dependent principle of neuromodulation: externally applied rhythmic stimulation preferentially amplifies the neural processes that actively support ongoing cognition, and does so in proportion to their recruitment. This principle provides a unified explanation for why stimulation effects are both task- and region-specific, and how identical stimulation parameters yield distinct neurophysiological changes under different cognitive demands. Specifically, when a neural process is strongly recruited by a task, it exhibits more substantial stimulation-induced modulation; when engagement is low, stimulation effects are correspondingly diminished. This principle held consistently across distinct oscillatory measures, i.e., local theta power and inter-regional theta synchrony, and across both WM tasks, underscoring its generality as an organizing rule for stimulation-brain interactions.

The engagement-dependent principle has significant scientific and translational implications. For example, stimulation protocols may need to be tailored to individuals based on their neural engagement profiles, including personalized task difficulty, stimulation timing, frequency, and targeted regions (Bueren et al., 2021; Fan et al., 2024; Soleimani et al., 2023). Moreover, rather than concentrating on presumed “poor performance states”, closed-loop interventions may be more effective when delivered during periods of strong engagement of the targeted mechanism, including during optimal performance (Bergmann, 2018; Sitaram et al., 2017). Mechanistically, our results help clarify the variability of stimulation outcomes reported in the neuromodulation literature (Guerra et al., 2017; Homan et al., 2021): variation in the engagement of the targeted mechanism, rather than inconsistency in stimulation parameters, may be a key determinant of stimulation efficacy.

The selective emergence of frequency-matching benefits only during the sequence WM task, but not the item WM task, provides further support for the engagement-dependent principle. While the entrainment hypothesis predicts that tACS efficacy should increase when stimulation frequency aligns with an individual’s endogenous theta peak (Krause et al., 2022), this mechanism alone cannot explain why the correlation between frequency mismatch and dwPLI gain was absent in the item WM task. Instead, our findings indicate that frequency matching confers a measurable benefit only when the targeted neural process is actively engaged by the task. When the item WM task failed to engage this same oscillatory mechanism, even a perfect frequency match produced no measurable benefit. These findings do not refute frequency entrainment but rather place it within a broader framework: entrainment is most effective when the targeted neural process is already engaged by task demands. We therefore propose the engagement-dependent resonance model, which posits that tACS efficacy arises from a multiplicative interaction between (1) the alignment of stimulation frequency with endogenous oscillatory dynamics and (2) the active cognitive engagement of the relevant neural process. In this view, frequency entrainment provides the *potential* for modulation, but task engagement serves as the *gate* that realizes this potential.

While the primary aim of the present study was to establish the engagement-dependent principle, our results also reinforce and extend prior work on item and sequence WM. Consistent with previous evidence, item WM relies on local theta power modulations (Brookes et al., 2011; Jensen & Tesche, 2002; Maurer et al., 2015), with frontal theta oscillations underpin the prioritization of task-relevant WM content (de Vries et al., 2018; Riddle et al., 2020). In contrast, sequence WM depends on theta phase coupling across distributed brain regions to maintain ordered representations (Johnson et al., 2018; Su et al., 2024; Yang et al., 2021), with frontoparietal theta phase synchronization increases with memory load and cognitive demand (Payne & Kounios, 2009; Sauseng et al., 2005; Schack et al., 2005)—a pattern indicative of executive control recruitment (Mizuhara & Yamaguchi, 2007). Inter-regional theta phase synchronization is further enhanced during tasks requiring integration of objects with temporal or spatial contexts (Benchenane et al., 2010; Jones & Wilson, 2005; Kay, 2005). Building on the seminal work of Polanía et al. (2012), which demonstrated that in-phase bifocal tACS enhances WM by promoting frontoparietal theta synchronization, the present study addresses why and under what conditions such stimulation is effective. By directly comparing two WM tasks with dissociable theta mechanisms, we show that identical in-phase stimulation produces task-specific effects, and that the magnitude of neural modulation scales with task engagement. These findings provide a mechanistic account of variability in tACS outcomes and introduce the engagement-dependent principle as a unifying framework.

Several methodological choices strengthened our conclusions. First, our design used identical stimuli, timing, and stimulation parameters across both tasks, thereby eliminating alternative explanations. Peripheral sensations (e.g., tingling, phosphenes) and E-field differences between stimulation conditions were matched across tasks and thus cannot account for the task-specific effects. Consequently, the only remaining factor driving the differential tACS effects between sequence and item WM tasks is the task requirement itself. Second, individualizing task difficulty across participants allowed more precise estimation of neural engagement. Third, MEG provided high temporal and spatial resolution for characterizing oscillatory dynamics and their modulation by tACS, which is crucial for examining region-specific oscillatory mechanisms. Nevertheless, we acknowledge that our MEG investigation focused on the bifocal condition, precluding direct examination of unifocal theta tACS effects. Still, our sample underrepresented male participants (approximately 26% male), although the observed effects were consistent across the full sample. Future studies should use these same design principles to explore whether engagement-dependent modulation generalizes to other frequency bands, cognitive domains, imaging modalities (e.g., functional magnetic resonance imaging [fMRI] and intracranial electroencephalography [iEEG]), and clinical populations. Computational models could further elucidate how external electrical fields interact with task-engaged neural circuits (Caul et al., 2020; Wischnewski et al., 2021).

In conclusion, our findings demonstrate that the effects of tACS on neural activity and behavior depend not only on stimulation parameters but also critically on the task-engaged neural mechanisms at the time of stimulation. The engagement-dependent principle reconciles task- and region-specific findings in the neuromodulation literature and provides a mechanistic foundation for designing more precise and effective stimulation strategies aimed at enhancing cognition.

## Method and Materials

### Participants

A total of 88 healthy right-handed college students (23 males; age range: 18–28 years) were recruited for this study. All participants had normal or corrected-to-normal visual acuity and no history of neurological or psychiatric disorders. Written informed consent was obtained from each participant after the experimental procedures were fully explained. Among the recruited participants, 39 completed the first behavioral experiment (Experiment 1), 26 finished the second behavioral experiment (Experiment 2), and 23 participated in the third MEG experiment (Experiment 3). No adverse effects related to tACS were reported during or after the experiments. The study protocol was approved by the Ethics Committee of the State Key Laboratory of Cognitive Neuroscience and Learning, Beijing Normal University (202009-29).

### Stimuli and procedures

#### Overview

Three experiments were designed to investigate the behavioral and neural effects of tACS on item and sequence WM (Fig. 1A). Experiment 1 spanned four days: one pre-screening day and three tACS days. Experiments 2 and 3 each included three days (one pre-screening day and two tACS days), with Experiment 3 additionally incorporating post-stimulation MEG scanning. The pre-screening day and the first tACS day were conducted consecutively, with subsequent tACS sessions spaced at least two days apart to avoid carryover effects. The pre-screening phase involved task familiarization (∼10 min) and memory capacity assessment (∼50 min) across four memory tasks. Each tACS day included task performance during (∼20 min) and after (∼40 min) stimulation.

In Experiment 1, participants were randomly divided into two groups: one group (N=19, 4 males, age range: 18–24 years) received unifocal tACS (6 Hz, 35 Hz, or sham) targeted at the left prefrontal cortex (F3) across three days; the other group (N=20, 4 males, age range: 19–24 years) received bifocal tACS (6 Hz in-phase, 6 Hz anti-phase, or sham) targeted at the left frontoparietal network (F3-P3). Experiments 2 (N=26, 5 males, age range: 18–24 years) and 3 (N=23, 10 males, age range: 20–28 years) exclusively employed bifocal 6 Hz in-phase and anti-phase tACS over two intervention days. Participants were randomly assigned to experimental groups, with task order and stimulation conditions counterbalanced across individuals but consistent for each participant throughout the study.

### Experiment tasks

Two task types (item WM and sequence WM) were applied. Both task types utilized two stimulus categories—fractal images (Mandelbrot Fractal Image Set) and spatial locations—to ensure that the dissociable effects of tACS on item and sequence WM generalize across qualitatively different representational domains, rather than being confined to stimulus-specific processing pathways. This yielded four experimental tasks: Fractal Image Memory, Fractal Image Sequence Memory, Spatial Memory, and Spatial Sequence Memory. Fractal Image tasks used abstract fractal images, while spatial tasks presented targets within a predefined 8×10 grid (screen resolution: 1920×1080) with gray squares randomly assigned to grid cells (Fig. 1B for detailed specifications).

#### Fractal Image Memory Task

Each trial began with a 0.5-s central fixation cross, followed by sequential presentation of high-quality fractal images (500 ms per stimulus). After a 2-s delay period (during which participants maintained fixation to memorize stimuli), a 4×4 matrix of 16 alternative images was displayed. Participants identified previously presented targets via mouse (Experiment 1) or keyboard (Experiments 2–3) without order constraints. A 1-s fixation cross marked the end of each trial.

#### Fractal Image Sequence Memory Task

Trials initiated with a 0.5-s fixation cross, followed by sequential presentation of fractal images (500 ms per item). After a 2-s memorization interval, stimuli reappeared in randomized order. Participants reconstructed the original sequence using mouse (Experiment 1) or keyboard (Experiments 2–3), with a 1-s intertrial interval.

#### Spatial Memory Task

Following initial fixation, an 8×10 grid with gray squares was displayed for 1 s. After a 2-s delay, participants recalled target locations via mouse/keyboard selection. Trials concluded with a 1-s blank screen.

#### Spatial Sequence Memory Task

An occluded 8×10 grid sequentially displayed gray squares (300 ms per item). After a 2-s delay, participants reconstructed the sequence via mouse/keyboard input, with a 1-s post-trial interval.

Due to MEG scanner constraints (only non-magnetic devices are permitted), Experiment 3 adopted numeric keyboard responses for all tasks (Fig. S1). To verify that response mode changes did not alter results relative to Experiment 1, Experiment 2 was conducted in a behavioral room using identical numeric keyboard inputs. For the Identity Memory Task’s probe phase, 10 alternatives were arranged in a 2×5 matrix and labeled 0–9. For Spatial Memory Tasks, each target location was sequentially probed with 10 surrounding alternatives (also numbered 0–9). Both sequence WM tasks (identity/spatial) used randomized numeric labeling. Participants in Experiments 2–3 responded sequentially: the left hand pressed keys 1–5, and the right hand pressed keys 6–0.

### Experimental procedures

#### Pre-screening Day

Participants first completed task training to familiarize themselves with the four WM tasks, followed by the memory capacity assessment phase. During capacity assessment, participants sequentially completed all four tasks with 1-minute breaks between tasks. Initial task loads were set at minimum levels: 1 item for Fractal Image and Spatial Memory Tasks, and 2 items for Fractal Image Sequence and Spatial Sequence Memory Tasks. Capacity assessment followed an adaptive one-up-two-down rule: task load increased by 1 after a correct response and decreased by 1 after two consecutive errors. Each task included 50 trials (∼12.5 minutes). Individual memory capacities for each task were calculated post-assessment (see “Memory Capacity Detection Analysis” section).

#### tACS Stimulation Days

Participants received 20 minutes of concurrent tACS while performing the four WM tasks. The on-stimulation phase included 20 trials per task (∼5 minutes), followed by a 1-minute break before the post-stimulation phase (40 trials (∼10 minutes) for Experiments 1 & 2; 50 trials (∼12.5 minutes) for Experiment 3). Task loads were set at each participant’s pre-determined capacity plus 1 to detect potential tACS-induced memory improvements.

### Memory capacity detection analysis

Following completion of all memory capacity assessment tasks on the pre-screening day, individual memory capacities were computed for each task using the following three-step process: first, identification of the maximum task load reached during the adaptive assessment phase; second, calculation of the average accuracy across all task loads from the initial load to the identified maximum load; finally, summation of these average accuracies to derive the preliminary capacity value.

For sequence WM tasks (Fractal Image Sequence and Spatial Sequence Memory), the initial load was set to 2, so the final capacity was determined by adding 1 to the summed average accuracies. For item WM tasks (Fractal Image and Spatial Memory), the preliminary summed average accuracies directly served as the final capacity (no additional adjustment was required).

Table S1 summarizes the memory capacities of all participants (N=88) across the three experiments. Since these capacity values were used to set task loads for subsequent tACS days (which required integer values), a rounding rule was applied: fractional parts ≥0.75 were rounded up, while fractional parts < 0.75 were truncated. For example, a summed average accuracy of 3.78 for the Identity Memory Task yielded a final capacity of 4, whereas a value of 3.23 was truncated to 3.

### Transcranial Alternating Current Stimulation (tACS)

Electrical stimulation was delivered via a battery-powered stimulator (NeuroConn GmbH, Ilmenau, Germany) with two or three 5×5 cm rubber electrodes placed on the scalp.

Prior to electrode placement, the scalp was prepared with conductive gel to ensure impedance levels remained below 10 kΩ. In Experiment 1, unifocal stimulation was administered with electrodes positioned over F3 (active) and C1 (return) according to the International 10-20 system, delivering sinusoidal waveforms (6 Hz or 35 Hz) at 2 mA peak-to-peak amplitude without DC offset. For bifocal stimulation in Experiments 1–3, electrodes were placed over F3 and P3 (active sites). The in-phase protocol involved theta (6 Hz) tACS with a 0° phase difference, applying 1 mA to F3 and P3 and 2 mA to C1 (return); the anti-phase protocol used theta (6 Hz) tACS with a 180° phase difference, delivering 1 mA to both F3 and P3. To visualize cortical E field distributions induced by each stimulation protocol, we performed finite-element method (FEM) simulations using SimNIBS v4.0 (Thielscher et al., 2015) on a standard head model (MNI-152 Colin27) with five tissue types: scalp, skull, cerebrospinal fluid, gray matter, and white matter. Electrodes were modeled as 5 × 5 cm rubber pads, 2 mm thick, positioned according to the 10–20 system (F3, P3, C1). Conductivities were assigned as follows: scalp 0.465 S/m, skull 0.010 S/m, CSF 1.654 S/m, gray matter 0.276 S/m, white matter 0.126 S/m (Opitz et al., 2015). E field magnitude was computed as the norm of the electric field vector. Simulations were performed for each protocol, and resulting E field distributions were visualized on the cortical surface (Fig. 1C). All active stimulations lasted 20 minutes, including 30-second ramping periods at the onset and offset. Sham stimulation involved only a 30-second ramp-up phase before immediate cessation (no sustained stimulation).

### MEG recordings and pre-processing

Post-stimulation tasks across the two tACS days in Experiment 3 were performed in the MEG scanner at Peking University. MEG data were acquired using a whole-brain Elekta Neuromag TRIUX system (Elekta, Stockholm, Sweden), equipped with 102 magnetometers and 204 planar gradiometers. Data were recorded at a sampling rate of 1000 Hz with an online band-pass filter (0.1–330 Hz). For accurate co-registration with MRI coordinates, three anatomical landmarks (nasion, left and right pre-auricular points), four head position indicator (HPI) coils, and at least 200 scalp and facial points were digitized using the Probe Position Identification system (Polhemus, Colchester, VT, USA). Structural MRI scans were acquired for all participants using a 3T Siemens Prisma scanner (Siemens Healthineers, Erlangen, Germany) with a 3D T1-weighted MPRAGE sequence (field of view [FOV] = 256 × 256 mm, repetition time [TR]/echo time [TE]/flip angle [θ] = 2530 ms/2.98 ms/7°, slice thickness = 1 mm, matrix = 256 × 256).

Raw MEG data were processed using MaxFilter software (Elekta-Neuromag) to apply temporal signal space separation (Taulu & Simola, 2006) for external noise removal.

Offline preprocessing of MEG data was conducted using the FieldTrip toolbox (http://www.fieldtriptoolbox.org) (Oostenveld et al., 2011) implemented in MATLAB (MathWorks Inc., Natick, MA, USA). The preprocessing pipeline included: (1) semi-automatic detection and exclusion of data segments with strong jumps or muscle artifacts; (2) application of a 50 Hz notch filter to remove line noise and its harmonics; (3) high-pass filtering at 1 Hz to eliminate slow drifts; (4) downsampling from 1000 Hz to 100 Hz to optimize processing efficiency and signal quality; (5) independent component analysis (ICA) for artifact removal (e.g., eye movements, head movements); (6) final visual inspection to ensure data quality. All subsequent analyses were performed using this preprocessed data.

### Time-frequency power analysis

The preprocessed MEG data were initially segmented into 6-second epochs spanning from 2 seconds before to 4 seconds after the final stimulus offset, designed to capture neural activity during the memory maintenance phase. This extended epoch duration was specifically chosen to mitigate edge artifacts introduced by subsequent time-frequency transformations. Analyses focused exclusively on a 2.5-second window centered on stimulus offset (0.5 s pre-offset to 2 s post-offset), with peripheral segments systematically discarded. Spectral decomposition was performed using complex Morlet wavelets with dual parameterization: 3–6 cycle kernels across 30 linearly spaced frequency bins (1–30 Hz) and 6–12 cycle kernels across 70 logarithmically spaced frequency bins (31–100 Hz). The resultant spectral power values were z-transformed for each channel-frequency pair, using trial-specific means and standard deviations calculated per experimental condition.

### Inter-regional phase synchronization analysis

The debiased weighted phase-lag index (dwPLI) was employed to quantify phase synchronization between brain regions (e.g., frontal-parietal lobes) within the target theta frequency band (Vinck et al., 2011). This method evaluates phase consistency across time points and trials using the following equation:

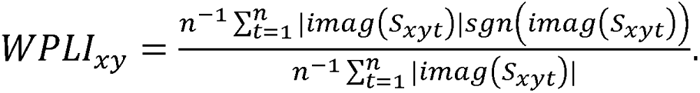

The dwPLI is robust against volume conduction artifacts, making it well-suited for functional connectivity analysis, with higher values indicating stronger inter-regional phase synchronization. For each trial, dwPLI values were computed for all 306 pairwise channel combinations (Vinck et al., 2011), with channels categorized into eight bilateral brain regions (occipital, temporal, parietal, and frontal lobes). Regional connectivity was assessed by averaging dwPLI values across 36 edges, including 28 inter-regional and 8 intra-regional connections. Analyses focused exclusively on the theta band (5–7 Hz) during the memory maintenance phase (0–2 s post-stimulus offset).

### Estimation of task-engaged theta peak frequency

To estimate each participant’s task-engaged endogenous theta peak frequency, we used MEG data from the anti-phase stimulation condition, which provides a measure of ongoing oscillatory activity without tACS-induced synchronization (Fig. 4C). For each participant and each task (item WM, sequence WM), MEG data were segmented into trials time-locked to the memory maintenance period (0–2 s post-stimulus offset). A Fast Fourier Transform (FFT) with a Hanning window was applied to the concatenated trials for each channel, and power spectra were computed within the theta band (4–8 Hz) at 0.1 Hz resolution. For each channel, the peak frequency was extracted as the frequency with maximum power. These peak frequencies were then averaged across channels within the left frontal and left parietal regions separately. The participant-specific task-state theta peak frequency was defined as the frequency with the maximum power averaged across these two regions. Finally, the mismatch between the fixed stimulation frequency (6 Hz) and the individual peak frequency was computed as:

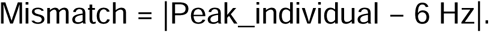

### Statistical analysis

#### Behavioral performance

To investigate the behavioral effects of tACS on memory performance in Experiment 1, post-stimulation accuracy was computed for item and sequence WM tasks across stimulation conditions. Three-way ANOVAs were conducted with WM task (item vs. sequence), stimulus (fractal vs. spatial), and stimulation condition as factors: for unifocal tACS, stimulation conditions included 6 Hz, 35 Hz, and sham; for bifocal tACS, conditions were in-phase, anti-phase, and sham. Experiments 2 and 3 exclusively employed 6 Hz in-phase and anti-phase stimulation. On-stimulation task accuracy was excluded due to insufficient trial counts. To assess group-specific tACS effects, behavioral gain metrics were derived (unifocal: ACC_6Hz_ – ACC_35Hz_; bifocal: ACC_in-phase_ – ACC_anti-phase_), followed by a mixed-design ANOVA with WM task, stimulus material, and stimulation group (between-subjects) as factors.

#### Non-parametric cluster-based permutation test

The non-parametric cluster-based permutation test (Maris & Oostenveld, 2007) was applied to identify clusters of time-frequency power exhibiting reliable differences between remembered and forgotten trials. This procedure sums neighboring t-values exceeding a cluster-forming threshold and compares the resulting cluster sizes to a null distribution of maximal cluster sums derived from Monte-Carlo label swapping.

Specifically, the empirical cluster t-value (remembered vs. forgotten) was calculated by summing neighboring t-values above a two-sided alpha threshold (0.025), with neighboring channels defined via FieldTrip’s triangulation method (minimum cluster size: 3 channels). The cluster-level t-value represented the sum of all within-cluster t-values, which was compared against a null distribution of 1000 surrogate t-values generated by random label permutations (two-sided alpha: 0.025). Corrected significance was determined by ranking the empirical t-value against the surrogate distribution.

#### Increases in inter-regional phase synchronization

First, three-way ANOVAs (WM task: item vs. sequence × stimulus: fractal vs. spatial × SME: remembered vs. forgotten) were conducted on average dwPLI values (from anti-phase stimulation trials) to identify edges exhibiting significant SMEs. Subsequently, three-way ANOVAs (WM task × stimulus × stimulation condition: in-phase vs. anti-phase) were performed on average dwPLI values (from all in-phase or anti-phase trials) to determine edges significantly modulated by tACS. Brain-behavior associations were examined by analyzing correlations between dwPLI gains (dwPLI_in-phase_ – dwPLI_anti-phase_) and behavioral gains (ACC_in-phase_ – ACC_anti-phase_) for both item and sequence WM tasks.

Finally, Cohen’s d effect sizes were calculated for the SME and tACS effect on each individual edge, and correlations between these effect sizes were examined separately for item and sequence WM tasks.

## Acknowledgements

This work was supported by the National Natural Science Foundation of China (32330039, 32441110 and 82271494), 111 project (BP071903), Beijing Natural Science Foundation (L256007), and Basic Research Program of Jiangsu (BK20250634).

## Author Contributions

Conceptualization, L.S. and G.X. Investigation, formal analysis, data curation, and visualization, L.S. Supervision, G.X. Writing – original draft, L.S. and G.X.

## Data availability

Data supporting this study are available on https://www.scidb.cn/en/anonymous/emllWVJq.

