## Supplementary materials for "Engagement-Dependent Neural Entrainment Underlies Dissociable tACS Effects on Item and Sequence Working Memory"

Table S1 Memory capacity for all four memory tasks of all participants (N=88)

| Experiments | Participants ID | Identity | Identity sequence | Spatial | Spatial sequence |
| --- | --- | --- | --- | --- | --- |
| Experiment 1 | 1 | 2.612 | 4.708 | 4.704 | 5.192 |
|  | 2 | 2.417 | 2.567 | 3.038 | 4.205 |
|  | 3 | 3.37 | 2.806 | 3.21 | 3.937 |
|  | 4 | 2.682 | 2.395 | 2.876 | 3.821 |
|  | 5 | 1.465 | 1.883 | 1.942 | 3.153 |
|  | 6 | 2.844 | 2.679 | 3.99 | 3.36 |
|  | 7 | 2.429 | 3.643 | 1.919 | 4.013 |
|  | 8 | 2.664 | 3.486 | 2.381 | 4.308 |
|  | 9 | 3.665 | 2.5 | 2.558 | 4.153 |
|  | 10 | 2 | 2 | 1.5 | 3 |
|  | 11 | 1.99 | 2.602 | 1.941 | 3.474 |
|  | 12 | 2.133 | 2.55 | 1.893 | 3.94 |
|  | 13 | 1.591 | 1.907 | 2.042 | 4.078 |
|  | 14 | 2.15 | 2.11 | 2.806 | 4.808 |
|  | 15 | 2.386 | 2.589 | 2.237 | 4.767 |
|  | 16 | 1.524 | 2.585 | 2.591 | 4.568 |
|  | 17 | 1.808 | 2.148 | 3.044 | 3.588 |
|  | 18 | 1.95 | 2.464 | 3.072 | 4.045 |
|  | 19 | 2.086 | 2.535 | 1.896 | 4.139 |
|  | 20 | 2.467 | 3.095 | 2.699 | 3.567 |
|  | 21 | 2.239 | 3.367 | 2.569 | 4.193 |
|  | 22 | 2.356 | 2.72 | 2.539 | 3.728 |
|  | 23 | 2.092 | 2.789 | 3.189 | 3.666 |
|  | 24 | 1.478 | 1.984 | 2.086 | 3.917 |
|  | 25 | 2.771 | 3.872 | 3.687 | 3.847 |
|  | 26 | 2.675 | 2.405 | 2.188 | 3.567 |
|  | 27 | 1.999 | 2.813 | 2.276 | 2.543 |
|  | 28 | 1.988 | 2.295 | 1.558 | 3.667 |
|  | 29 | 1.846 | 3.488 | 3.022 | 4.304 |
|  | 30 | 2.041 | 2.721 | 3.727 | 3.814 |
|  | 31 | 1.966 | 2.688 | 2.653 | 4.556 |
|  | 32 | 2.304 | 2.647 | 2.335 | 3.3 |
|  | 33 | 1.881 | 2.874 | 2.277 | 4.082 |
|  | 34 | 2.475 | 2.761 | 2.17 | 3.885 |
|  | 35 | 2.139 | 3.514 | 1.764 | 4.343 |
|  | 36 | 2.058 | 3.017 | 3.397 | 4.069 |
|  | 37 | 2.242 | 2.513 | 2.723 | 3.872 |
|  | 38 | 1.63 | 2.421 | 2.128 | 3.395 |
|  | 39 | 2.371 | 3.739 | 3.149 | 3.79 |
| Experiment 2 | 40 | 3.102 | 4.508 | 2.933 | 4.697 |
|  | 41 | 3.233 | 2.458 | 1.178 | 4.522 |
|  | 42 | 2.328 | 4.045 | 2.276 | 5.833 |
|  | 43 | 5.008 | 6.01 | 1.977 | 5.712 |
|  | 44 | 2.047 | 3.04 | 2.123 | 4.176 |
|  | 45 | 3.286 | 4.771 | 1.952 | 4.875 |
|  | 46 | 3.937 | 3.75 | 2.106 | 4.571 |
|  | 47 | 2.182 | 3.27 | 1.545 | 4.123 |
|  | 48 | 3.429 | 5.917 | 1.69 | 4.666 |
|  | 49 | 2.298 | 2 | 1.231 | 4.471 |
|  | 50 | 1.742 | 4.071 | 2.438 | 4.611 |
|  | 51 | 3.481 | 3.988 | 2.04 | 4.869 |
|  | 52 | 2.27 | 3.971 | 2.355 | 4.261 |
|  | 53 | 3.094 | 4.98 | 1.617 | 4.093 |
|  | 54 | 3.2 | 4.574 | 2.36 | 4.713 |
|  | 55 | 3.571 | 5.627 | 1.929 | 4.273 |
|  | 56 | 3.326 | 4.299 | 3.3 | 4.893 |
|  | 57 | 2.466 | 4.973 | 1.637 | 4.387 |
|  | 58 | 2.537 | 2.995 | 2.148 | 4.377 |
|  | 59 | 2.642 | 4.884 | 1.408 | 4.33 |
|  | 60 | 2.432 | 2.739 | 1.719 | 3.519 |
|  | 61 | 3.086 | 6.085 | 2.098 | 4.628 |
|  | 62 | 3.231 | 3.959 | 2.21 | 5.221 |
|  | 63 | 3.243 | 2.194 | 1.417 | 4.152 |
|  | 64 | 1.111 | 2.289 | 1.25 | 4.053 |
|  | 65 | 1.111 | 2.289 | 1.25 | 4.053 |
| Experiment 3 | 66 | 2.889 | 4.076 | 1.188 | 4.089 |
|  | 67 | 3.089 | 3.611 | 2.142 | 3.937 |
|  | 68 | 1.979 | 2.393 | 2.432 | 4.819 |
|  | 69 | 3.678 | 4.542 | 1.959 | 5.11 |
|  | 70 | 3.658 | 2.705 | 1.699 | 5.033 |
|  | 71 | 2.709 | 4.668 | 1.39 | 3.815 |
|  | 72 | 5.512 | 5.611 | 2.176 | 5.407 |
|  | 73 | 1.968 | 3.064 | 1.744 | 4.034 |
|  | 74 | 4.024 | 4.382 | 1.994 | 4.574 |
|  | 75 | 1.574 | 2.853 | 1.46 | 4.18 |
|  | 76 | 3.422 | 3.229 | 2.114 | 5.001 |
|  | 77 | 3.122 | 4.221 | 2.068 | 4.613 |
|  | 78 | 3.721 | 4.832 | 2.475 | 5.561 |
|  | 79 | 3.019 | 4.037 | 2.06 | 4.808 |
|  | 80 | 3.361 | 4.722 | 2.368 | 4.433 |
|  | 81 | 2.689 | 3.739 | 2.296 | 5.293 |
|  | 82 | 2.5 | 4.8 | 2.344 | 4.828 |
|  | 83 | 2.22 | 4.743 | 1.921 | 4.902 |
|  | 84 | 3.56 | 4.579 | 1.867 | 4.767 |
|  | 85 | 3.816 | 4.029 | 2.658 | 5.171 |
|  | 86 | 3.6 | 3.838 | 1.867 | 4.91 |
|  | 87 | 2.621 | 5.085 | 2.013 | 5.268 |
|  | 88 | 2.839 | 4.633 | 3.881 | 4.748 |

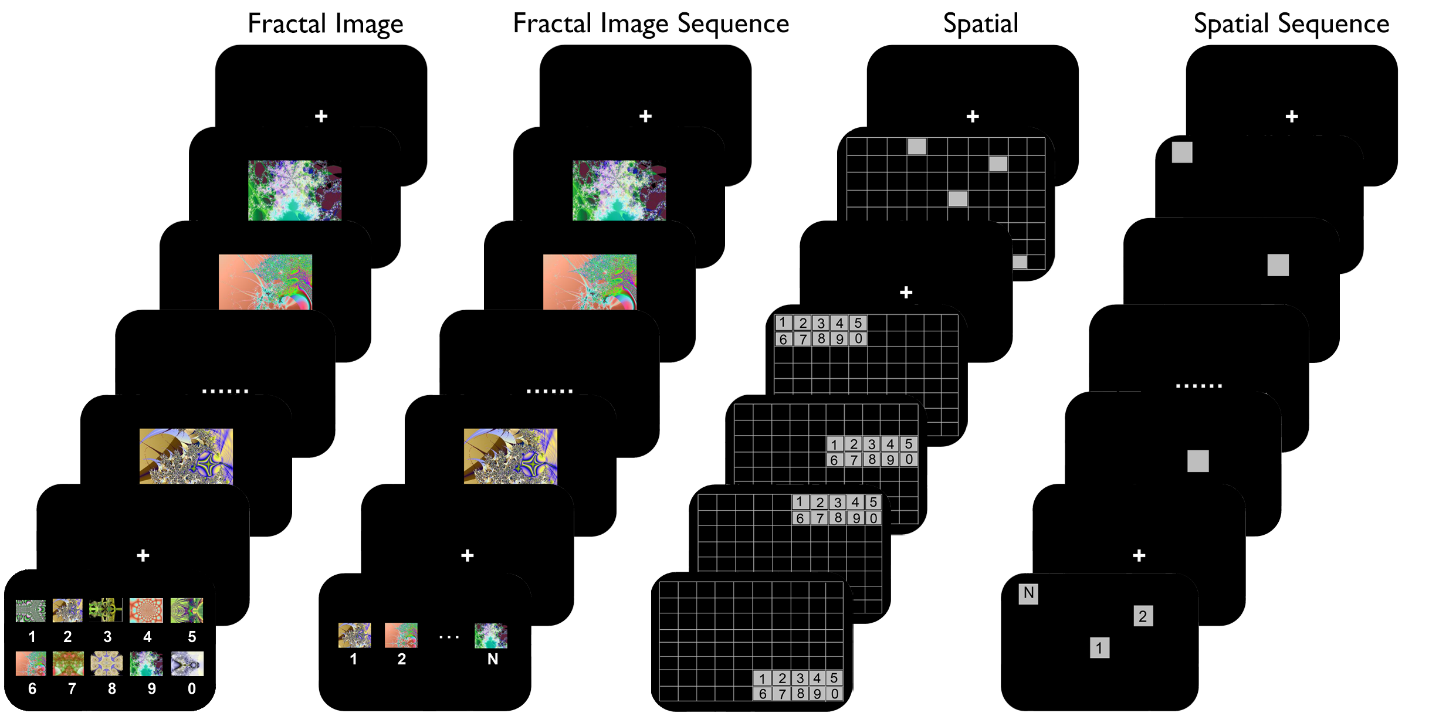

**Fig. S1.** **Schematic representation of the item working memory and sequence working memory paradigms in the Experiments 2 and 3**. All settings and parameters in Experiments 2 and 3 were identical to those in Experiment 1, except for the response mode, which was changed from mouse to keyboard. For the probe phase of the Fractal Image Memory Task, 10 alternative fractal images were presented in a 2×5 grid labeled with keys 0–9. For the Spatial Memory Task’s probe phase, each target location was probed sequentially, with 10 surrounding alternative locations labeled 0–9. In contrast, for both the Fractal Image Sequence and Spatial Sequence Memory Tasks, numerical labels (0–9) were randomly assigned to stimuli to guide keyboard-based sequence reconstruction.
